# Music-evoked reactivation during continuous perception is associated with enhanced subsequent recall of naturalistic events

**DOI:** 10.1101/2025.07.05.663273

**Authors:** Jamal A. Williams, Elizabeth H. Margulis, Christopher Baldassano, Uri Hasson, Janice Chen, Kenneth A. Norman

## Abstract

Music is a potent cue for recalling personal experiences, yet the neural basis of music-evoked memory remains elusive. We address this question by using the full-length film Eternal Sunshine of the Spotless Mind to examine how repeated musical themes reactivate previously encoded events in cortex and shape next-day recall. Participants in an fMRI study viewed either the original film (with repeated musical themes) or a no-music version. By comparing neural activity patterns between these groups, we found that music-evoked reactivation of neural patterns linked to earlier scenes in the default mode network was associated with improved subsequent recall. This relationship was specific to the music condition and persisted when we controlled for a proxy measure of initial encoding strength (spatial intersubject correlation), suggesting that music-evoked reactivation may play a role in making event memories stick that is distinct from what happens at initial encoding.

## 1 Introduction

Music alters and enriches our lives, shaping our experiences and coloring our memories. Imagine attending a party where many unfamiliar songs were played. Several days following the party, you are driving and one of the songs plays on the radio. Suddenly, you are transported back to the party, remembering the people who attended, conversations you had, and even the decorations that embellished the house. Accordingly, numerous laboratory studies have demonstrated that music is a powerful retrieval cue (Smith, 1985; Boltz et al., 1991; Janata et al., 2007; Janata, 2009; Belfi et al., 2016; Kubit & Janata, 2022). Here, we investigate how music-cued memory reactivation relates to subsequent recall of naturalistic events; in the above example, how does remembering details from the party in response to the repeated song relate to your ability to remember these details later on? Numerous studies have shown that the act of retrieving a memory is a particularly potent way of strengthening that memory (Roediger and Butler, 2011; Liu et al., 2012; Antony et al., 2017), suggesting that the musical reminder should help to make the details from the party “stick” in your mind, but existing work has not addressed this question directly.

Our specific hypothesis was that stronger reactivation of stored event memories (in response to repeated musical cues) would be associated with better subsequent recall of these events. To test this hypothesis, it is necessary to first *measure reactivation*, by recording the neural pattern at encoding and assessing how strongly that pattern is evoked by the cue; having measured reactivation, one can assess whether higher levels of reactivation are associated with better subsequent recall. While there have been multiple fMRI studies (Janata, 2009; Ford et al., 2011; Falcon et al., 2022) looking at how music can evoke autobiographical memories (so-called “music-evoked autobiographical memories”, or MEAMs), none of them have measured neural reactivation: The standard approach to studying MEAMs is to present experimenter-selected music from various periods of participants’ lives and measure (through self-report) retrieval of idiosyncratic autobiographical memories. Because the evoked memories in these studies were formed outside of the lab (and therefore outside of the scanner), these studies were not able to measure reactivation of neural patterns from encoding.

As in these studies of MEAMs, we wanted to explore how music can trigger recall of rich naturalistic content, but – in our case – we wanted to bring the encoding process under experimental control so we could measure reactivation and relate it to subsequent recall. To accomplish this goal, we scanned 24 participants with fMRI while they watched the full-length film *Eternal Sunshine of the Spotless Mind*. We chose this film because its soundtrack contains six recurring musical themes that are repeated a variable number of times throughout the film. This allowed us to collect neural data when a musical theme was first heard during a scene and also during subsequent re-exposure to that musical theme, when it could serve as a reminder of the initial scene. We also scanned 24 participants in a *no-music control* condition – this condition was exactly the same as the main condition, except that the music soundtrack was removed from the film, keeping all other sounds (i.e. dialogue, ambient sounds) intact (see *Methods*). The day after viewing the movie, participants from both groups returned to perform a surprise spoken recall test for the entire movie.

Using this experimental design, we measured the extent to which repetition of musical themes during movie-watching triggered neural reactivation of earlier scenes that had been paired with those themes; we predicted that higher levels of music-evoked reactivation of earlier scenes would be associated with better subsequent recall of these scenes. Crucially, we expected that reactivation in response to repeated themes would go beyond activation of the neural pattern for the song (something we would expect from an area involved in musical perception, even if it had no involvement in memory) to also encompass reactivation of non-musical aspects of earlier scenes; this fits with the idea, from the initial “party” example, that the repeated music should also trigger retrieval of what happened at the party, what the party looked like, and so on. Based on prior work showing that brain regions in the default mode network (DMN) represent the contents of naturalistic events (e.g., Chen et al., 2016; Chen et al., 2017; Zadbood et al., 2017; Lee et al., 2021; Yeshurun et al., 2021; Barnett et al., 2024), we predicted that these reactivation effects would be localized to the DMN (i.e., that greater reactivation of scene-specific event representations in the DMN would be associated with improved subsequent memory for those scenes).

## 2 Results

### 2.1 Music-evoked reactivation in DMN areas is associated with enhanced recall

Figure 1a shows how we measured music-evoked memory reactivation. A key challenge with using pattern similarity to track memory reinstatement is that high levels of pattern similarity between earlier and later scenes could be driven simply by *perception of features that are shared* between the two scenes, as opposed to *retrieving a memory* of the earlier scene. In our study, if we were to directly compare neural patterns evoked by two scenes where participants were listening to the same musical theme, the similarity might be driven simply by perception of that musical theme, as opposed to retrieval of non-musical details. To address this challenge, we took advantage of our unique two-group design and compared the similarity of later scenes from the *music* group to earlier scenes from the *no-music* group. Comparing across groups allowed us to rule out the possibility that pattern similarity was being driven by perception of the same musical theme; rather, any similarity that we observed would have to reflect non-musical features of the earlier scene. Specifically, reactivation was measured in two steps: First, we computed the pattern similarity between a) each music participant’s neural activity for scenes containing a given song and b) the average of the no-music group’s neural activity for *earlier scenes containing the same song* (green arrow in Figure 1a). This similarity score can reflect both memory retrieval of features of the earlier scene (evoked by the repeated theme) and also any perceptual similarity between the earlier and later scenes. To control for this latter factor, we subtracted out the pattern similarity of those scenes in the no-music group (red arrow in Figure 1a). This procedure yielded a reactivation score for each scene containing a (subsequently) repeated musical theme for each music participant (see *Methods* for details; note that this approach of comparing scene patterns across participants is supported by numerous prior naturalistic fMRI studies that have found shared scene-specific information across participants; e.g., Chen et al., 2017; Zadbood et al., 2017). Next, for each participant in the music condition, we sorted their scene reactivation scores based on whether scenes were subsequently remembered or subsequently forgotten, and then averaged the scores within each bin, resulting in a remembered reactivation score and a forgotten reactivation score for each participant (Figure 1b). We then compared the remembered reactivation scores to the forgotten reactivation scores using one-tailed paired samples t-tests (remembered > forgotten).

**Figure 1.**
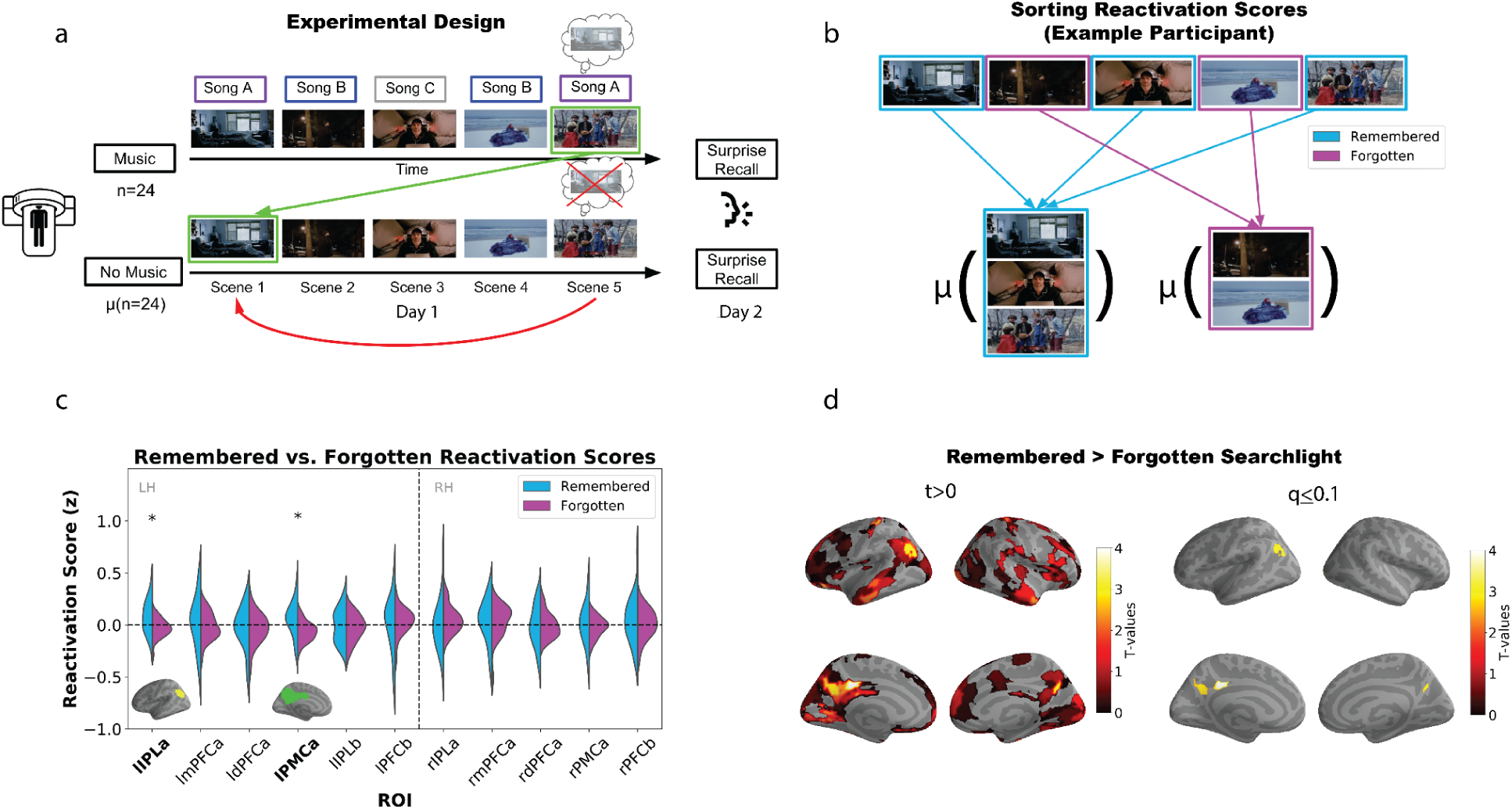
(a) We tested for reactivation of non-musical information in the music participants by comparing their event patterns for later scenes containing a particular song to the average of the no-music group’s previous scenes containing the same song (green arrow), while controlling for baseline pattern similarity between scenes (red arrow). Participants performed a free recall test the next day, where they were asked to recall the entire movie in as much detail as possible. (b) For each participant, reactivation scores were computed for each scene, then sorted into remembered or forgotten bins and averaged within each bin. (c) Remembered and forgotten reactivation scores for pre-defined DMN ROI clusters were compared using t-tests. Significant differences between remembered and forgotten reactivation scores were observed in left PMC (*t*(23) = 3.047, q < 0.05) and left IPL (*t*(23) = 2.614, q < 0.05). (d) **Left**. Parcel-based searchlight (remembered > forgotten) t-test results (uncorrected map thresholded at t > 0). **Right**. FDR-corrected t-test results within the full DMN (DMNa, DMNb, and DMNc). Regions in left and right posterior medial cortex and left angular gyrus survive our threshold criterion (q ≤ 0.1); all of these regions fall within the DMNa subnetwork.

Initially, we restricted our analysis to the default mode network (DMN), dividing it into the DMNa and DMNb subnetworks using the Schaefer atlas (Schaefer et al., 2018). Recent evidence suggests that DMNa is involved in episodic memory, whereas DMNb has been linked to social behaviors (DiNicola et al., 2020). Based on these findings, we hypothesized that reactivation scores would be most strongly related to subsequent recall (i.e., the remembered vs. forgotten difference would be largest) in DMNa.

We observed that reactivation scores for remembered scenes were significantly higher than those for forgotten scenes in the left inferior parietal lobule (IPLa; *t*(23) = 2.614, q < 0.05) and the left posterior medial cortex (PMCa; *t*(23) = 3.047, q < 0.05; Figure 1c). Additionally, we performed a similar parcel-based searchlight analysis across the full brain (Figure 1d, left panel); since our *a priori* hypothesis was that this effect would be localized to the DMN, we corrected for multiple comparisons within the DMN (including DMNa, DMNb, and DMNc). This analysis confirmed higher reactivation scores for remembered scenes in left IPLa and bilateral PMCa, again demonstrating specificity to the DMNa subnetwork (Figure 1d, right panel; for statistics for these parcels see Table S5).

In addition to looking at the relationship between reactivation and subsequent recall behavior, we also analyzed these two measures separately. When we tested for reactivation (without sorting into bins of remembered and forgotten scores) within pre-defined DMN ROIs, we did not observe significant reactivation in these regions (Supplementary Figure 4a). However, when we performed the analysis as a parcel-based searchlight, we found a significant main effect of reactivation in right angular gyrus (*t*(23) = 3.957, q < 0.05) and left parahippocampal cortex (*t*(23) = 2.968, q < 0.1; Supplementary Figure 4b). When looking at behavior on its own, we did not find a significant difference in the amount of scenes recalled between the music and no-music conditions (Supplementary Figure 3a); however, we did find that – when we directly compared recall of particular scenes in the music and no music conditions (limiting ourselves to scenes where music was played in the music condition) – the proportion of participants who recalled a given scene tended to be higher in the music condition than the no-music condition (Supplementary Figure 3b). We address the relationship between these various findings in the *Discussion* section.

### 2.2 Analyses controlling for initial encoding strength, measured using intersubject correlation (ISC)

There are two possible interpretations of the observed relationship between music-evoked reactivation and subsequent recall: One is that reactivation causes improved subsequent recall; the other possibility is that reactivation and subsequent recall are indirectly related via a third variable, the *strength of encoding of the earlier scene*. According to this latter view, strong encoding of the earlier scene leads to higher levels of reactivation of that scene (when the musical theme is repeated) and also better subsequent recall of the earlier scene on the final recall test, inducing a correlation between reactivation and subsequent recall; importantly, this relationship can hold even if there is no direct causal relationship between reactivation and subsequent recall. We set out to tease these two possibilities apart by first acquiring a measure of initial encoding strength, then regressing that measure from the reactivation scores, and finally observing whether the relationship between reactivation and subsequent recall is still present. If the relationship between reactivation and subsequent recall is solely due to initial encoding strength, then regressing it out will eliminate this relationship. However, if the effect persists, then we can conclude that it is not solely due to initial encoding strength.

For this analysis, we used spatial intersubject correlation (ISC) as our proxy measure of initial encoding strength. Spatial ISC measures the similarity of the spatial patterns evoked by a particular scene across participants (Nastase et al., 2019). Intuitively, when a participant is encoding a scene attentively, their neural representation of the scene will resemble that of other participants, but when a participant is inattentive / encoding poorly, their neural representation of the scene will diverge from that of other participants; in line with this intuition, prior work has shown that spatial ISC predicts subsequent memory (Meshulam et al., 2021; Koch et al., 2020), justifying its use here as a measure of initial encoding strength. For each brain parcel and for each scene containing a (subsequently) repeated theme, we computed the time-averaged spatial pattern for that scene a) in each individual participant and b) in the N-1 other participants, and we correlated those two patterns (Figure 2a), resulting in a spatial ISC score for each scene, in each brain parcel. We then measured whether these ISC scores differed for subsequently remembered vs. forgotten scenes. When we included results from both the music and no-music participants, we found that spatial ISC significantly predicted subsequent recall in the superior parietal lobe (q ≤ 0.1; Figure 2b), validating the use of ISC as a proxy for initial encoding strength in our study. In order to do the best possible job of predicting initial encoding quality *specifically in the music participants*, we identified the regions where ISC predicted subsequent recall at a liberal threshold of p < 0.05 (Figure 2c); 20 such regions were identified. Next, within each participant, we sequentially regressed out the ISC values for these 20 regions from the reactivation score for a given region. The purpose of this analysis was to remove any variance that was shared between the 20 regions’ ISC scores (proxies of initial encoding strength) and reactivation; we opted to use a liberal threshold for including a region’s ISC score in this analysis because we wanted to err on the side of removing more (vs. less) variance in encoding strength. The final residuals (i.e., variance in reactivation not explained by ISC across any of the ROIs) were then used to predict subsequent recall.

**Figure 2.**
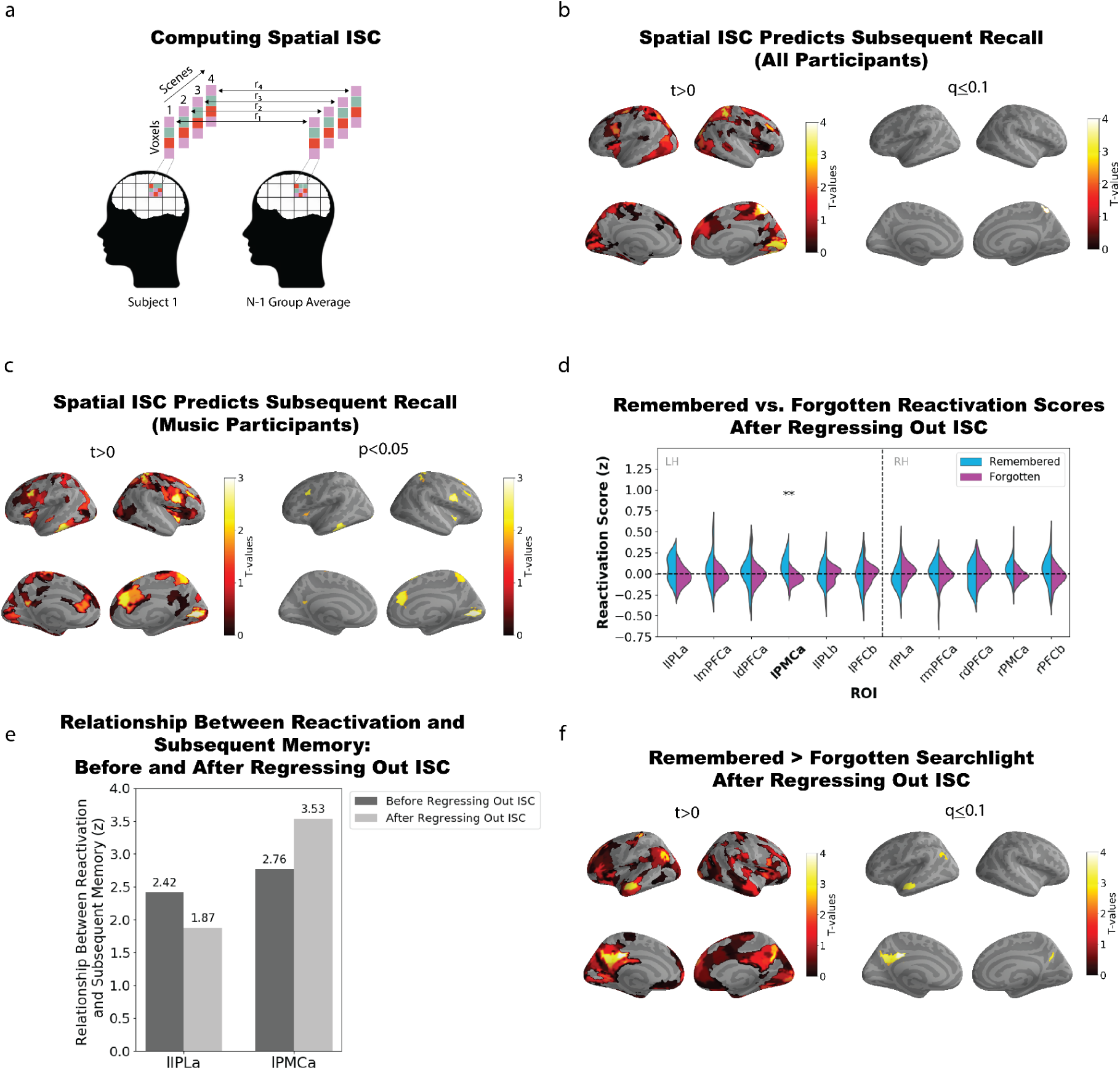
(a) Calculation of spatial intersubject correlation (ISC). (b) Regions showing a relationship between ISC and subsequent recall when participants in both conditions (music and no-music) are included in the analysis. **Left**. Regions with t-values greater than zero. **Right**. Regions passing false discovery rate correction (q ≤ 0.1); right superior parietal lobe exhibits a significant relationship between ISC and subsequent recall. (c) Regions showing a relationship between ISC and subsequent recall, when we limit the analysis to music participants. **Left**. Regions with t-values greater than zero. **Right**. Regions where ISC predicts subsequent recall at a liberal threshold of p < 0.05, uncorrected for multiple comparisons. (d) Reactivation scores after regressing out ISC (see text) were compared for subsequently remembered and forgotten scenes, in pre-defined DMN ROI clusters. A significant difference between remembered and forgotten reactivation scores was observed in left posterior medial cortex (*t*(23) = 4.122, q < 0.005). (e) Comparison of relationship between reactivation and subsequent recall before regressing out ISC (dark gray) and after regressing out ISC (light gray) in left IPL and left PMC (the two regions that showed a significant effect in Figure 1c). (f) Brain map from parcel-based searchlight analysis showing parcels where there was a significant relationship between reactivation scores (after regressing out ISC) and subsequent recall. **Left**. Regions with t-values greater than zero. **Right**. Regions passing false discovery rate correction (q ≤ 0.1).

When using these residual reactivation scores to predict subsequent recall (in the same *a priori* ROIs used in Figure 1c above), we found that PMC remained significant (q < 0.005) whereas IPL did not (Figure 2d). Figure 2e directly compares the strength of the relationship between reactivation and subsequent recall, pre and post regression, showing that the IPL effect numerically drops whereas the PMC effect numerically increases. Figure 2f shows the relationship between reactivation (controlling for ISC using the above regression method) and subsequent recall using a parcel-based searchlight approach. Significant effects passing false discovery rate correction were observed in one parcel in left angular gyrus, five parcels in left PMC, one parcel in right PMC, and also one parcel in the left temporal lobe – an effect not present in the original (pre-regression) version of the analysis; for statistics for these parcels see Table S6. Overall, the fact that the relationship between reactivation and subsequent recall survived (or even increased numerically) in several regions after regressing out ISC suggests that this relationship is not an artifact of initial encoding strength driving both reactivation and subsequent recall (we return to this issue in the *Discussion*).

### 2.3 No evidence of reactivation in the no-music condition

Next, we performed a control analysis to confirm that the relationship between reactivation and subsequent recall (described above) is only observed in the presence of music. Our primary reactivation analysis controls for the *average* level of non-musical similarity between scenes (the red arrow in Figure 1a); however, it does not control for the possibility that idiosyncratic differences in attention to non-musical features could drive these effects. For example, if a participant happens to attend especially strongly to the main character’s journal that is also present in an earlier scene, this could trigger increased memory reactivation of the earlier scene and (through this) improved subsequent recall of the scene, leading to a relationship between increased reactivation and subsequent recall, even in the absence of music; subtracting out the average level of perceptual similarity would not compensate for these idiosyncratic effects.

To address this possibility, we repeated the initial subsequent recall analysis, but – instead of comparing music participants’ later scenes to no-music participants’ earlier scenes – we compared held-out no-music participants’ later scenes to other no-music participants’ earlier scenes. No significant differences between remembered and forgotten reactivation scores were observed using the pre-defined ROIs (Figure 3a) or the parcel-based searchlight procedure (Figure 3b).

**Figure 3.**
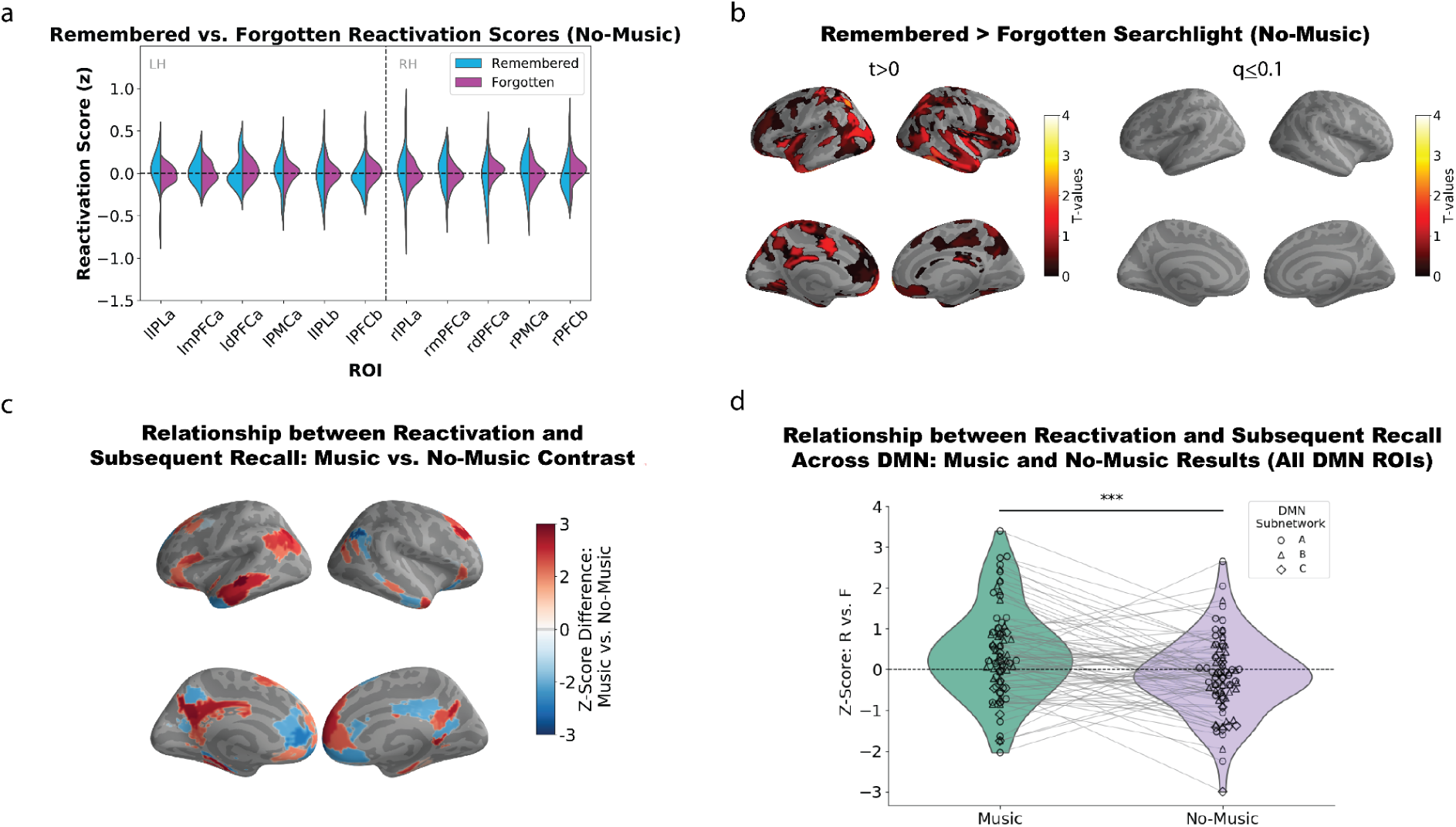
(a) We performed a control analysis to test whether there was a relationship between reactivation and subsequent recall in the no-music group. No significant differences were observed between remembered and forgotten reactivation scores in our predefined ROIs (belonging to DMNa and DMNb) for the no-music participants. (b) **Left**. Parcel-based searchlight (remembered > forgotten) t-test results. **Right**. FDR-corrected t-test results within the full DMN (DMNa, DMNb, and DMNc). Searchlight analysis revealed no significant evidence of a relationship between reactivation and subsequent recall in the DMN. (c) Analysis comparing remembered vs. forgotten differences in reactivation (expressed as z-scores) between the music and no-music conditions within the full DMN. Red regions represent areas where the remembered vs. forgotten difference was larger for the music condition, and blue regions represent areas where the remembered vs. forgotten difference was larger for the no-music condition. (d) Violin plots showing remembered vs. forgotten z-scores for all 79 DMN parcels, as a function of condition. Across DMN parcels, remembered vs. forgotten differences in reactivation were larger in the music condition than the no-music condition (*t*(78) = 3.92, p < 0.0001). Parcels belonging to different DMN subnetworks are visualized using different shapes (see legend).

Next, to test whether the subsequent recall effect in the DMN was greater in the music condition than the no-music condition, we first computed the difference in the size of the subsequent recall effect for music vs. no music participants separately for each DMN parcel (Figure 3c; see also Supplementary Figure 2); then, we assessed whether this difference was reliable across parcels using a one-tailed paired samples t-test (music > no music) with degrees of freedom equal to the number of DMN parcels minus one (see *Methods* for details). Here, we observed a significant difference between the music and no-music conditions: Across the 79 DMN parcels, the size of the subsequent recall effect was consistently higher in the music group than the no-music group (*t*(78) = 3.92, p < 0.0001; Figure 3d). Note that, while this analysis approach lets us assess reliability across DMN parcels, it treats participants as a fixed effect and therefore does not license generalization to new participants. As an alternative to this approach, we tried another analysis variant where we computed the reliability of the remembered vs. forgotten difference across DMN parcels separately for each participant, and then we assessed whether this reliability value was larger in music vs. no-music participants (see *Methods* for details). This analysis gets at a similar question to the previous analysis but licenses generalization to new participants; here, we found a marginal difference between the music and no-music participants, *t*(46) = 1.57, p = 0.06.

### 2.4 Music-evoked reactivation is specific to the DMN and visual areas

Finally, we sought to determine whether music-evoked reactivation was specific to the DMN. We re-ran our analysis comparing the size of the subsequent recall effect between the music and no-music conditions (the first variant of the analysis, with degrees of freedom equal to the number of ROIs minus one), this time looking at parcels in each of 9 bilateral cortical networks generated from the Schaefer 400 parcellation atlas (see *Methods*). We found that, in addition to the DMN showing a relationship between reactivation and subsequent recall, this relationship was also present in visual areas. Specifically, the visual peripheral network also showed a significant difference between the music and no-music conditions (*t*(22) = 2.93, q = 0.017; Figure 4). No other networks showed significant evidence of a relationship between reactivation and subsequent recall.

**Figure 4.**
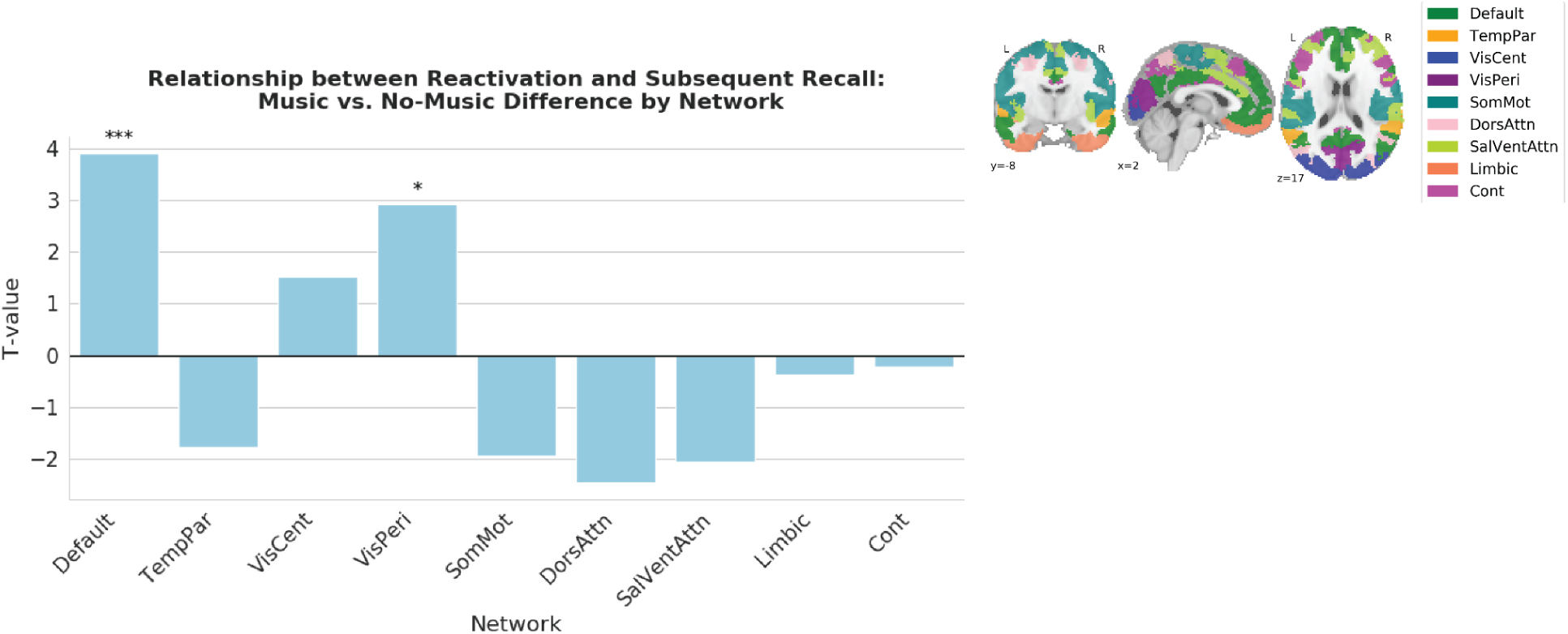
Plot showing the difference in the relationship between reactivation and subsequent recall for music vs. no-music participants, separately for each cortical network. Each bar represents a t-value acquired by comparing the size of the subsequent recall effect (difference in R and F reactivation, expressed as a z-score) between the music and no-music conditions, across all ROIs within each cortical network. In addition to the DMN, the visual peripheral network exhibits a greater subsequent recall effect in the music condition compared to the no-music condition (*t*(22) = 2.93, q = 0.017).

## 3 Discussion

In this study, our main prediction was that repeated musical themes would reactivate the DMN representations of previous events associated with those themes, and that this reactivation would be associated with improved subsequent memory. In keeping with this prediction, we showed that music-evoked reactivation of *non-musical* scene details was higher for remembered than forgotten scenes in posterior medial cortex and angular gyrus. This relationship between reactivation and subsequent recall could be attributable to reactivation causing better subsequent recall, but it could also be attributable to both of these factors being driven by a third variable, initial encoding strength. To arbitrate between these possibilities, we used spatial intersubject correlation (ISC) as a proxy for initial encoding strength, first showing that ISC predicts subsequent recall and then regressing it out from reactivation. We found that, while the relationship between reactivation and subsequent recall was numerically diminished in some areas when we regressed out ISC, the relationship was still significant (and even numerically larger) in many areas, including posterior medial cortex; this suggests that reactivation may have effects on subsequent recall that go beyond the effects of initial encoding strength (see discussion below). We further demonstrated that, across regions in the DMN, this relationship between reactivation and subsequent recall was larger in participants who heard the musical themes than in a control group who viewed a version of the movie without the musical themes. Finally, in an exploratory analysis, we showed that the DMN and the visual peripheral network were the only brain networks that show this association between music-evoked reactivation and subsequent recall. Our finding that music-evoked reactivation of narrative events occurs in regions of the DMN fits with recent work showing that the DMN represents high-level event structure in both audiovisual narratives (Chen et al., 2017; Baldassano et al., 2017; Baldassano et al., 2018; Song et al., 2021; Yeshurun et al., 2021; Grall et al., 2023; Reagh & Ranganath, 2023) and music (Williams et al., 2022). As mentioned in the *Introduction*, a limitation of prior fMRI studies that have used musical cues to elicit autobiographical memories (Janata, 2009; Ford et al., 2011; Falcon et al., 2022) is that – in those studies – there was no objective way to measure the strength of memory reactivation, since the memories being retrieved were encoded outside of the scanner. Our study builds on this prior work by providing an objective way of measuring music-evoked memory retrieval, and by demonstrating that this measure relates to subsequent recall.

Our unique two-group design, where some participants viewed the movie with the music intact and others viewed the movie with the music removed, contributed to our conclusions in two important ways. First, by comparing later *music* scenes to earlier scenes from the *no-music* group (see Figure 1), we could rule out the possibility that pattern similarity was being driven by perception of the same musical theme. Second, by running our reactivation analysis separately in the music and no-music groups, we were able to show that residual *non-musical* similarity between earlier and later scenes was not sufficient to drive the effect – similarity in non-musical features was exactly matched between the two participant groups, but the relationship between pattern similarity and subsequent recall was significantly larger across the DMN (and the visual peripheral network) in the participants who heard the musical themes. Taken together, the results provide strong evidence that our key effect was driven by reinstatement of non-musical features, triggered by the repeated musical cues.

More generally, our study contributes to a growing literature showing that accurate memory retrieval is a powerful experience that can improve the subsequent accessibility of the retrieved memory (Tulving, 1967; Carrier & Pashler, 1992; Butler, 2010; Rowland, 2014; Schuetze et al., 2019; for a review, see Roediger & Abel, 2022). Much of this literature has focused on measuring how explicitly testing someone’s memory affects subsequent recall (the so-called “testing effect”; Roediger & Karpicke, 2006; Roediger & Butler, 2011). Our results suggest that incidental reminders during continuous perception of naturalistic experience may strengthen memories for previously perceived events; i.e., it is not necessary to pause presentation of the movie stimulus and ask someone to intentionally retrieve in order to observe these memory benefits (but see caveats below). Our work resembles studies of *targeted memory reactivation* (TMR; Antony et al., 2012; Tambini et al., 2017; Wang et al., 2019; Hu et al., 2020; Nicolas et al., 2025; for reviews, see Oudiette & Paller, 2013, Antony & Paller, 2017, and Carbone & Diekelmann, 2024) in some respects. TMR studies, like ours, look at how incidental presentation of retrieval cues (i.e., without instructions to retrieve) affects subsequent memory. However, unlike our study, TMR studies typically present retrieval cues by themselves (i.e., they pause presentation of new stimuli) and have used static stimuli (e.g., objects presented on a grid) as to-be-remembered materials (e.g., Rasch et al., 2007; Rudoy et al., 2009; Antony et al., 2018), instead of looking at memory for dynamic naturalistic events, although a few have looked at memory for dynamic naturalistic events using virtual reality (Shimizu et al., 2018; Borghese et al., 2022).

As noted above, a limitation of our study is that the observed relationship between reactivation and subsequent recall need not be causal; the relationship could instead be attributable to the third variable of initial encoding strength. The results of our control analysis in Section 2.2 (showing that the observed relationship between reactivation and subsequent recall was numerically *larger* in some regions after regressing out ISC) suggest that variance in initial encoding strength is not responsible for the observed relationship in those regions, but this analysis is not definitive. ISC was the only measure of initial encoding strength that we had access to in this study; while ISC is clearly related to subsequent recall, it does not capture all of the variance in initial encoding strength, and it is possible that regressing out some other measure of initial encoding strength (not available to us here) could have led to different results. Future work can more definitively address this causality question by experimentally manipulating reactivation within participants; this could be accomplished, e.g., by keeping repeated themes for some scenes but removing the repeated themes for other scenes, and then measuring whether recall is higher for scenes where the repeated themes are kept vs. removed.

Our work also builds on other fMRI studies that have looked at “reminding” within naturalistic stimuli. For example, Cohn-Sheehy et al. (2021) found increased pattern similarity in the hippocampus for “coherent” narrative events that were part of the same plot, compared to unrelated events; Chang et al. (2021) found increased pattern similarity in DMN areas for key narrative motifs that repeated across a story, compared to other repeated words that did not play a significant role in the narrative; Song et al. (2025) observed neural reactivation of causally-related past events at moments when participants reported sudden insight into narratives; and Hahamy et al. (2023) found increased reactivation at event boundaries of previous events that had high (vs. low) feature overlap with the just-completed event, in both the hippocampus and the DMN. Our study resembles these other studies in that we also used pattern similarity to track memory reactivation during narrative perception; the present work goes beyond these other studies in showing a relationship between cued reactivation strength for individual events and subsequent recall of these events. None of these other studies were set up to look for this relationship; the closest that these other studies came to our key result was the finding from Chang et al. (2021) that, across subjects, higher average levels of reinstatement triggered by key narrative motifs were associated with higher average levels of understanding of how earlier and later parts of the story (sharing those motifs) related to each other, as measured by a behavioral test (see also Song et al., 2025, for a similar result).

Additionally, our findings contribute to the understanding of music as a memory retrieval cue. Behavioral studies that have used music as a cue to retrieve autobiographical memories from earlier in life (acquired outside of the lab) suggest that musical cues can enhance retrieval (Janata, 2007; Belfi et al., 2016; Belfi & Jakubowski, 2021). However, results from controlled laboratory studies testing memory for in-lab events are more mixed. Most of these studies have arbitrarily paired music with simple stimuli (e.g., word lists, simple visual stimuli) and examined whether reinstating the music at retrieval enhances recall accuracy (measured behaviorally). While some of these studies have found a memory benefit when the same music is played at encoding and retrieval (Smith, 1985; Balch & Lewis, 1996; Mead & Ball, 2007), others failed to show this benefit or even showed a negative effect of music reinstatement in some conditions (e.g., Balch et al., 1992; Echaide et al., 2019). Another recent study paired short video clips with either unrelated music or other sound cues, and found that music elicited fewer correct memories compared to other sound cues, despite music-cued memories being rated as having a more positive valence (Jakubowski et al., 2023). One possible reason for these mixed results is that music tends to spontaneously prompt rich scene construction (as would be used when imagining narratives or reconstructing autobiographical events; Jakubowski et al., 2024); this could impair performance when music is used to cue retrieval of relatively meaningless events like individual words. Another possible reason for these mixed results is the arbitrary nature of the pairing between music and the to-be-remembered stimuli in the aforementioned studies: When music is not meaningfully related to ongoing events, it may act as a distraction rather than an effective retrieval cue. Indeed, studies have demonstrated that background music can impair performance during cognitive tasks (Salame & Baddeley, 1989; Kämpfe et al., 2011). Unlike prior studies, our paradigm featured music that was purpose-built by the filmmakers to be *congruent* with the events in the movie (e.g., the mood of the music that accompanies a scene typically aligns with the overall mood of that scene). The literature on schema-consistent learning (e.g., Audrain et al., 2022; Shin et al., 2021; McClelland, 2013; for a review, see Gilboa & Marlatte, 2017) suggests that this kind of congruency could facilitate the formation of associations between music and the events of the movie, making the music a more effective retrieval cue later on; we discuss ways of testing this hypothesis below.

Despite the purpose-built match between the music and the movie, the effects of music on memory in our study were still quite variable. There was a robust relationship between reactivation and subsequent recall in the DMN (Figures 1c, 1d, 2d, 2f, 3c, 3d, 4), but overall neural reactivation effects (not split by subsequent recall) were less consistent – they were not significant after multiple comparisons correction in any *a priori* ROIs (Supplementary Figure 4a) and only significant in two parcels in our parcel-based searchlight analysis (Supplementary Figure 4b) – and the main effect of music vs. no-music on behavioral recall performance was not significant in an unpaired t-test (Supplementary Figure 3a). Some insight into this pattern of results comes from our behavioral analysis where we compared the recall performance of music and no-music participants for individual scenes: While the music version of the scene tended (on average) to be recalled better than the no-music version of a scene, there was extensive variability across scenes in how strongly recall of that scene was benefited by music (Supplementary Figure 3b). The fact that recall of some scenes was not benefited by music (or potentially was even harmed by music) may explain the lack of a significant overall benefit of music on behavioral recall, and also why there were fewer regions showing an overall neural reactivation effect, compared to the number of regions showing a relationship between neural reactivation and subsequent recall.

Going forward, it will be important to address this question of what makes music a more effective cue in some circumstances than others. The most powerful way to address this will be through new studies that experimentally manipulate relevant factors. For example, we noted above that congruency between the music and the scene could be an important driver of reactivation effects in our study. The effects of congruency could be studied using a variant of the experiment that reassigns the themes across scenes while preserving the repetition structure (e.g., all of the scenes with theme A could be switched to theme B). It will also be useful to analyze how specific features of the musical stimuli affect their efficacy as retrieval cues. For example, a recent study showed that emotional intensity, pulse strength, brightness (strength of high harmonics), and frequency fluctuations (low-mid) are significant predictors of music-evoked autobiographical memories (Salakka et al., 2021). Our present study is not adequately powered to study these issues, but future work could address this by manipulating these features directly (e.g., adjusting the themes to reduce emotional intensity). Furthermore, while we have focused here on the efficacy of music as a *retrieval* cue, it is possible that the presence of music at encoding could also benefit recall (e.g., through the deployment of associative strategies; Ferreri et al., 2015) even if the music is never repeated later – indeed, our behavioral results show hints of this kind of effect (see Supplementary Figure 3 and associated discussion). Future work can address this by separately manipulating whether music is presented at encoding and whether it is repeated later.

Another question for future research is to explore the role of the hippocampus in these reactivation effects. Hippocampus was not one of the regions showing a significant relationship between reactivation and subsequent recall. While we need to be careful not to overinterpret null results, this fits with the view that detailed event content is primarily represented in neocortex, and hippocampus is responsible for storing *indices* that bind together these event-specific details, including any musical themes that accompany an event (McClelland et al., 1995; Teyler & DiScenna, 1986). According to this view, when musical themes are repeated, activity could spread from the neocortical representation of the musical theme into the hippocampus, activating the associated episodic memory (“index”) of the original event in the hippocampus; from here, activity can spread back out from hippocampus to the features of the original event in neocortex (“pattern completion”). Future work could explore hippocampal contributions, e.g., by using electrocorticography to measure hippocampus-to-cortex information transfer (Michelmann et al., 2021, 2025).

In conclusion, our study demonstrates that music-evoked reactivation of neural activity patterns in the default mode network and visual processing regions is associated with successful subsequent retrieval of naturalistic events. This relationship remained significant and was numerically enhanced in several regions after controlling for a proxy measure of initial encoding strength (spatial intersubject correlation), suggesting that reactivation may have effects on subsequent recall that are distinct from the effects of initial encoding strength. Additional work with new, purpose-built stimuli is needed to evaluate the (potentially) causal nature of this relationship and to identify which features of the music and scenes are most effective in driving music-evoked reactivation.

## Methods

### Participant sample

We collected fMRI data from a total of 48 participants (29 females, ages 18–36 years; Mean = 21.2 years, SD = 3.70 years). Participants were paid $20 per hour. 46 of the participants were native English speakers. The experimental protocol was approved by the institutional review board of Princeton University, and all participants gave their written informed consent.

### Experimental design

All participants watched the full-length film *Eternal Sunshine of the Spotless Mind* for a total duration of 1 hour and 48 minutes while undergoing fMRI scanning. Half of the participants watched the original version of the film with the musical soundtrack intact while the other half watched an edited version of the film with the musical soundtrack removed. A music-free version of the movie was created by purchasing the dialogue track (containing both dialogue and ambient sounds) and a 35mm audio-free version of the film from NBCUniversal Media, LLC. Next, we paired the audio-free film with the dialogue track, resulting in a music-free version of *Eternal Sunshine of the Spotless Mind*. Movie-viewing was split into two scanning runs so that participants could take a short break (maximum of 5 minutes). Movie-viewing resumed when participants stated they were ready to continue. All participants returned the following day for a surprise recall test in which they were asked to recall the entire movie in as much detail as possible while undergoing fMRI scanning. Total recall times ranged from 10 minutes to 66 minutes with an average recall time of 37 minutes and a standard deviation of 15 minutes (fMRI data from the recall phase are not reported in this paper). The experiment was implemented in Psychopy version 3.1.5.

### Scanning parameters and preprocessing

Results included in this manuscript come from preprocessing performed using fMRIPrep 1.2.3 (Esteban et al., 2018; Esteban et al., 2019; RRID:SCR_016216), which is based on Nipype 1.1.6-dev (Gorgolewski et al., 2011; Gorgolewski et al., 2018; RRID:SCR_002502).

### Anatomical data preprocessing

A total of 2 T1-weighted (T1w) images were found within the input BIDS dataset. All of them were corrected for intensity non-uniformity (INU) using N4BiasFieldCorrection (Tustison et al., 2010; ANTs 2.2.0). A T1w-reference map was computed after registration of 2 T1w images (after INU-correction) using mri_robust_template (FreeSurfer 6.0.1; Reuter et al., 2010). The T1w-reference was then skull-stripped using antsBrainExtraction.sh (ANTs 2.2.0), using OASIS as target template. Brain surfaces were reconstructed using recon-all (FreeSurfer 6.0.1, RRID:SCR_001847, Dale et al., 1999), and the brain mask estimated previously was refined with a custom variation of the method to reconcile ANTs-derived and FreeSurfer-derived segmentations of the cortical gray-matter of Mindboggle (RRID:SCR_002438, Klein et al., 2017). Spatial normalization to the ICBM 152 Nonlinear Asymmetrical template version 2009c (Yoon et al., 2009, RRID:SCR_008796) was performed through nonlinear registration with antsRegistration (ANTs 2.2.0, RRID:SCR_004757, Avants et al., 2008), using brain-extracted versions of both T1w volume and template. Brain tissue segmentation of cerebrospinal fluid (CSF), white-matter (WM), and gray-matter (GM) was performed on the brain-extracted T1w using fast (FSL 5.0.9, RRID:SCR_002823, Zhang et al., 2001).

### Functional data preprocessing

For each of the 3 BOLD runs that we collected per participant (across all tasks and sessions), the following preprocessing was performed. First, a reference volume and its skull-stripped version were generated using a custom methodology of fMRIPrep. A deformation field to correct for susceptibility distortions was estimated based on fMRIPrep’s fieldmap-less approach. The deformation field was that resulting from co-registering the BOLD reference to the same participant T1w-reference with its intensity inverted (Wang et al., 2017; Huntenburg, 2014).

Registration was performed with antsRegistration (ANTs 2.2.0), and the process was regularized by constraining deformation to be nonzero only along the phase-encoding direction, and modulated with an average fieldmap template (Treiber et al., 2016). Based on the estimated susceptibility distortion, an unwarped BOLD reference was calculated for a more accurate co-registration with the anatomical reference. The BOLD reference was then co-registered to the T1w reference using bbregister (FreeSurfer) which implements boundary-based registration (Greve & Fischl, 2009). Co-registration was configured with nine degrees of freedom to account for distortions remaining in the BOLD reference. Head-motion parameters with respect to the BOLD reference (transformation matrices, and six corresponding rotation and translation parameters) were estimated before any spatiotemporal filtering using MCFLIRT (FSL 5.0.9, Jenkinson et al., 2002). BOLD runs were slice-time corrected using 3dTshift from AFNI 20160207 (Cox & Hyde, 1997, RRID:SCR_005927). The BOLD time-series (including slice-timing correction when applied) were resampled onto their original, native space by applying a single, composite transform to correct for head-motion and susceptibility distortions. These resampled BOLD time-series will be referred to as preprocessed BOLD in original space, or just preprocessed BOLD. The BOLD time-series were resampled to MNI152NLin2009cAsym standard space, generating a preprocessed BOLD run in MNI152NLin2009cAsym space. First, a reference volume and its skull-stripped version were generated using a custom methodology of fMRIPrep. Several confounding time-series were calculated based on the preprocessed BOLD: framewise displacement (FD), DVARS and three region-wise global signals. FD and DVARS were calculated for each functional run, both using their implementations in Nipype (following the definitions by Power et al., 2014). The three global signals were extracted within the CSF, the WM, and the whole-brain masks. Additionally, a set of physiological regressors were extracted to allow for component-based noise correction (CompCor, Behzadi et al., 2007). Principal components were estimated after high-pass filtering the preprocessed BOLD time-series (using a discrete cosine filter with 128s cut-off) for the two CompCor variants: temporal (tCompCor) and anatomical (aCompCor). Six tCompCor components were then calculated from the top 5% variable voxels within a mask covering the subcortical regions. This subcortical mask was obtained by heavily eroding the brain mask, which ensures it does not include cortical GM regions. For aCompCor, six components were calculated within the intersection of the aforementioned mask and the union of CSF and WM masks calculated in T1w space, after their projection to the native space of each functional run (using the inverse BOLD-to-T1w transformation). The head-motion estimates calculated in the correction step were also placed within the corresponding confounds file. The BOLD timeseries were resampled to surfaces on the following spaces: fsaverage6. All resamplings can be performed with a single interpolation step by composing all the pertinent transformations (i.e. head-motion transform matrices, susceptibility distortion correction when available, and co-registrations to anatomical and template spaces). Gridded (volumetric) resamplings were performed using antsApplyTransforms (ANTs), configured with Lanczos interpolation to minimize the smoothing effects of other kernels (Lanczos, 1964). Non-gridded (surface) resamplings were performed using mri_vol2surf (FreeSurfer). Only movie encoding BOLD data were analyzed for this study.

### Statistical Analyses

#### Behavioral annotations

To relate reactivation to recall performance, we needed to establish scene descriptions that could be used to assess participants’ recall. Scenes were defined by a group of independent raters. First, a single rater watched the movie and segmented it into relatively short chunks based on whether they felt that a meaningful shift occurred between each one (Zacks & Swallow, 2007). These shifts could have pertained to shifts between entire scenes or important narrative transitions. Next, the same rater wrote short descriptions (a few sentences long) for each newly defined scene. Scene descriptions were provided at a high level (e.g. “Joel calls in sick to his boss at an empty, snow ridden train station, telling her he has food poisoning and goes to the beach.”). From the remaining group of raters (n=14), we obtained an additional set of event boundaries. This time, raters were asked to identify time points in the movie at which there were breaks in the flow: These could be scene changes, shifts of topic, location, time, action, emotion, or other factors that made them think that an event had ended and a new event had started.

Furthermore, raters were instructed that event boundaries could come close together or far apart and to try to identify between 4-20 boundaries for every 5 minutes of video. This was an approximate range and not an exact requirement. Finally, they were told to try to keep their criteria consistent across the entire movie. In this case, these raters only had to watch the movie and provide event boundaries and they did not have to provide event descriptions. This procedure was completed in two splits of the raters whereby one group watched the version of the film containing the music (n=8) while the other group watched the music-free version of the movie (n=6). Consensus boundaries were computed within each group as well as across both groups by smoothing over boundaries using a 3s Gaussian kernel and then thresholding the peaks at the 90th percentile. Given that our primary questions pertained to how music affects memory for event details, we decided to only use the consensus boundaries acquired from the group of raters who watched the version of the movie that contained music.

Next, we adjusted the scene boundaries provided by the first independent rater by comparing them to the consensus boundaries acquired from all other raters. For example, if two scenes defined by the first rater actually fell within a longer scene identified by the other raters, then these two scenes would now count as a single scene. The adjusted event boundaries were our final set of event boundaries for the film and each scene was then given a unique scene label. This resulted in 407 unique scenes.

Next, we needed to identify which scene a participant recalled by assigning a scene label (or scene labels) to each recall utterance made by the participant. Recall utterances were defined by having a group of independent raters (n=6) segment each participant’s recall transcript into individual sentences. Then, raters provided timestamps for each recall utterance by listening to where each utterance occurred in participants’ recall audio recordings. Each timestamp corresponded to the onset of the recall utterance in the audio recording. Finally, another group of independent raters (n=10; including 4 of the raters from the previous task) assigned scene labels to each recall utterance made by the participants (note that multiple labels could be assigned to a given utterance if the raters thought that the utterance reflected recall of multiple scenes). A scene was classified as “remembered” if at least one recall utterance was labeled with that scene.

#### Subsequent recall analysis

##### (1) Computing reactivation scores

**Identifying relevant scenes:** The movie featured 27 unique songs, of which 6 were repeated a variable number of times. Descriptive statistics for these 6 repeated songs were as follows:

Repetition frequency (in number of occurrences): mean = 2.16, median = 1.5, standard deviation = 1.46, min = 1, max = 5.

Song durations: mean = 70.32s, median = 59s, standard deviation = 44.09s, min = 12s, max = 173s.

Duration between repetitions: mean = 1316.54s, median = 642s, standard deviation = 1732.19s, min = 63s, max = 5405s.

Only the scenes associated with these 6 repeated songs were included in the analysis, resulting in 93 repeated music scenes distributed unequally across songs (Figure 5). Scenes per song ranged from 4 to 27 (mean = 15.5, standard deviation = 8.42). See *Supplementary Materials* for more information on the songs.

**Figure 5.**
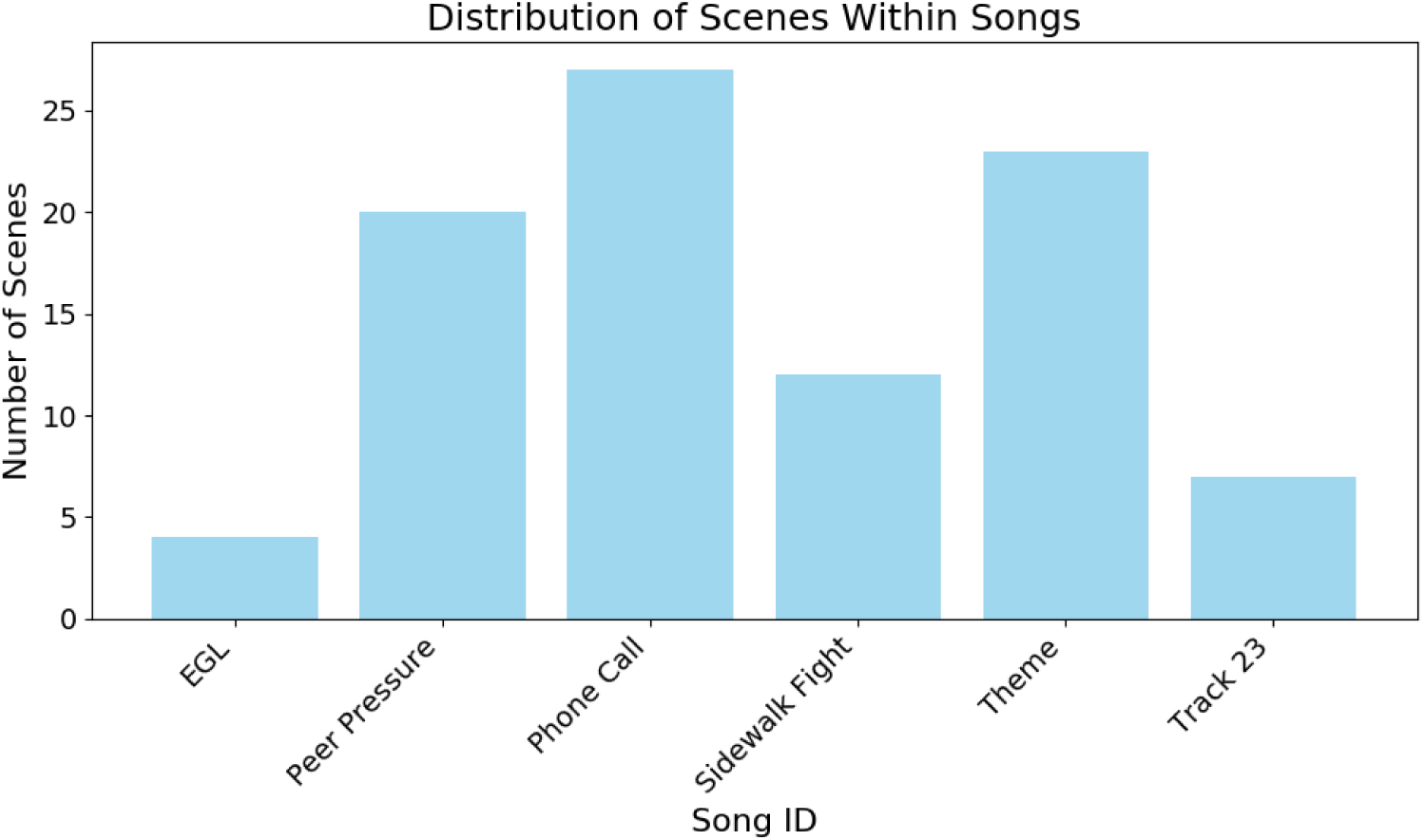
Number of scenes associated with each song. The total number of scenes containing repeated music was 93. Minimum number of scenes = 4 (EGL), max = 27 (Phone Call), mean = 15.5, SD = 8.42.

###### Extracting event patterns

To investigate whether non-musical information was reactivated by music, we extracted neural activity patterns for both the music and no-music conditions. For each participant, song-specific event patterns were computed by averaging neural activity across TRs within each of the 93 repeated music scenes. If the same song was playing across multiple successive scenes, the patterns for these scenes were averaged together into a single “mega-scene” (for example, the song “Phone Call” was played throughout scenes 111-115, so the patterns for these five scenes were averaged into a single mega-scene). For the no-music condition only: We averaged across no-music participants to create a single no-music group pattern for each of the 93 repeated music scenes.

###### Computing raw reactivation scores

Neural pattern similarity was computed between each music participant’s “mega-scenes” containing a given song and the no-music group’s earlier scenes containing the same song. In doing this, we made sure that the no-music group’s earlier scenes were *strictly earlier* than (i.e., not contained within) the mega-scene they were being compared to (e.g., in the “Phone Call” example, the music-group mega-scene composed of scenes 111-115 was only compared to the no-music group’s “Phone Call” scenes with indices < 111). We took this precaution of never comparing scenes within a “mega-scene” because pattern similarity within a mega-scene (e.g., comparing no-music 111 to music 115) could potentially reflect features held in working memory instead of retrieval from long-term memory. Our comparison procedure resulted in 67 correlation coefficients in total, referred to as the *raw reactivation scores*.

###### Controlling for baseline similarity

To control for baseline scene similarity, we calculated pattern similarity for each no-music participant using a leave-one-out approach. Specifically, we compared each no-music participant’s later mega-scenes to the group average’s earlier scenes, excluding that participant from the average. These baseline similarity scores were averaged across no-music participants and subtracted from the raw reactivation scores, resulting in *contrast reactivation scores* (see Figure 1a).

###### Creating null distributions

To account for chance-level similarity, we generated null distributions by randomly shuffling event patterns and recomputing the contrast reactivation scores 1000 times. The true scores were then converted to z-scores by subtracting the mean of the shuffled scores and dividing by their standard deviation. These final z-scores, simply referred to as *reactivation scores*, quantify the extent of music-evoked reactivation for each scene.

##### (2) Comparing remembered and forgotten reactivation scores

After computing reactivation scores for each scene containing repeated music, we sorted these scores based on whether the scenes were remembered or forgotten. This sorting resulted in two bins of reactivation scores per participant, with the number of scores in each bin varying across participants due to differences in the number of remembered scenes. Within each bin, reactivation scores were averaged, producing a single remembered reactivation score and a single forgotten reactivation score for each music participant. To assess whether reactivation was stronger for remembered scenes, we conducted a one-tailed independent samples t-test (remembered > forgotten).

We performed this analysis at two levels: ROI-based and parcel-based searchlight.

1. **ROI-level analysis**: For our ROI analysis, we targeted clusters within the default mode network (DMN) using the Schaefer 400 parcellation 17 network atlas (Schaefer et al., 2018). Specifically, we selected parcels from the DMNa and DMNb subnetworks to create custom ROIs for key regions. This approach was motivated by prior findings that implicate DMNa in episodic processing and DMNb in social behaviors (DiNicola et al., 2020):

○ Inferior Parietal Lobule (IPLa, IPLb)
○ Medial Prefrontal Cortex (mPFCa)
○ Dorsolateral Prefrontal Cortex (dlPFCa, dlPFCb)
○ Posterior Medial Cortex (PMCa) When possible, ROIs were divided by hemisphere (e.g., left and right), although some regions (e.g., IPLb) were only represented in one hemisphere in the Schaefer atlas. P-values for each ROI were corrected for multiple comparisons by implementing the false discovery rate (FDR) procedure used in AFNI.
2. **Parcel-based searchlight analysis**: For the searchlight analysis, we extended the remembered vs. forgotten comparison across all cortical parcels. For each parcel, we computed one-tailed t-tests comparing remembered and forgotten reactivation scores and assigned p-values; these p-values in the DMN were subsequently converted to q-values using FDR. The resulting map (Figure 1d) visualizes t-values thresholded at q ≤ 0.1, highlighting regions of the DMN where reactivation was significantly associated with subsequent recall.

#### Controlling for initial encoding strength

We wanted to assess the extent to which initial encoding strength could explain subsequent recall. Here we operationalize initial encoding strength as spatial intersubject correlation (ISC): the neural similarity between the time-averaged scene patterns for a given participant and the group average. We first determined the set of brain regions where ISC predicted subsequent recall. Then, we regressed these ISC scores from the reactivation scores to test whether reactivation still predicted subsequent recall. The analysis steps were as follows:

1. **Computing spatial ISC:** We conducted a spatial ISC subsequent recall analysis where, for each scene containing (subsequently) repeated music, we first correlated a given participant’s time-averaged scene pattern with the N-1 group average for that same scene. To allow for differences between music and non-music participants, ISC scores were computed within-condition (i.e., music participants were compared to the other music participants; no-music participants were compared to the other no-music participants). Next, scene-specific spatial ISC scores were sorted into bins based on whether they were remembered or forgotten, and then these ISC scores were averaged within each bin for each participant. This resulted in an average remembered and forgotten spatial ISC score for each participant within each condition. We then performed a one-tailed (remembered > forgotten) paired samples t-test; this was done once including participants from both conditions (music and no-music), and once including only music participants. These analyses were performed as a parcel-based searchlight using the Schaefer 400 parcellation atlas. Regions where ISC predicted subsequent recall in the music condition at the liberal threshold of p < 0.05 uncorrected were selected for further analyses; there were 20 regions that passed this threshold.
2. **Regressing out spatial ISC:** We repeated the analysis relating reactivation scores to subsequent recall, but here we first regressed out 20 spatial ISC values (corresponding to the 20 regions passing the p < .05 criterion noted above) from each participant’s reactivation scores before relating the residuals to subsequent recall. This regression was done in 20 successive steps. In each step, one region’s ISC values were used as a predictor in a linear model, with the current residuals from the previous step as the response. The residuals were updated after each regression, resulting in a final residual vector that captured variance in reactivation not explained by ISC across any of the 20 selected regions. These residuals were then binned according to whether the corresponding scenes were subsequently remembered or forgotten, and averaged within each bin. All subsequent steps followed the same procedure as the original subsequent recall reactivation analysis.

#### Running the subsequent recall analysis in the no-music condition

As a control analysis, we wanted to assess the degree to which reactivation related to subsequent recall in the no-music participants. Performing this analysis would help us further determine whether the reactivation observed in the music participants was actually related to the music. To this end, we repeated the subsequent recall analysis, except – instead of comparing music participants’ later mega-scenes to no-music participants’ earlier scenes – we compared held-out no-music participants’ later mega-scenes to other no-music participants’ earlier scenes. Specifically, we computed raw reactivation scores by comparing each no-music participant’s later mega-scenes to the other no-music participants’ earlier scenes containing the same song (averaging across these other no-music participants to get a single pattern for each of the earlier scenes). Furthermore, we also excluded the no-music participant used to compute the raw reactivation score from the calculation of the corresponding baseline reactivation score. All other steps of the analysis were the same as the original subsequent recall analysis.

#### Comparing subsequent recall effects between music and no-music conditions in the DMN

To evaluate whether the subsequent recall effect on reactivation was greater in the music condition than in the no-music condition in the DMN, we tried two different analysis approaches.

For the first analysis variant, we first computed the difference in the size of the subsequent recall effect for music vs. no music participants separately for each DMN parcel, and then we assessed whether this was reliable across parcels. Specifically, for each of the 79 DMN parcels across the DMNa, DMNb, and DMNc networks, we ran a one-tailed paired samples t-test (remembered > forgotten reactivation) across participants, separately for the music and no-music participants; we then normalized the resulting test statistics by converting them to z-scores (Supplementary Figure 2). Next, we calculated the difference in z-scores for each DMN parcel between the two conditions (music – no-music); these difference scores indicate the degree to which the music-condition subsequent recall effect exceeded that of the no-music condition. We plotted these differences on the surface for visual inspection (Figure 3c). Finally, to measure the statistical significance of the difference between conditions, we performed a one-tailed paired samples t-test (music > no-music) on the per-parcel z-scores (Figure 3d), with degrees of freedom equal to the number of DMN parcels minus one.

For the second analysis variant, we computed the reliability of the remembered vs. forgotten difference across DMN parcels separately for each participant, and then we assessed whether this reliability value was larger in music vs. no-music participants. Specifically, for each participant, we computed a remembered reactivation score and a forgotten reactivation score for each DMN parcel. Next, within each participant, we ran a one-tailed paired samples t-test (remembered > forgotten reactivation) to measure the reliability of the remembered vs. forgotten reactivation difference across DMN parcels (with degrees of freedom equal to the number of DMN parcels minus one). Next, we converted the per-participant t-values to z-scores, and we did a one-tailed, two-sample t-test on these z-scores comparing the music and no-music groups.

#### Computing subsequent recall effects between music and no-music conditions across all networks

To assess whether the observed difference in subsequent recall between the two conditions was specific to the DMN, we extended the above analysis (specifically, the first variant of the analysis) to all other networks in the Schaefer atlas. This analysis was performed across all parcels within each network by collapsing across hemispheres and subnetworks, resulting in 9 distinct networks derived from the Schaefer 400 parcellation 17 network atlas. For each network, we calculated subsequent recall z-scores for each parcel in that network and conducted a one-tailed paired samples t-test (music > no-music) on the per-parcel z-scores. This produced a t-value for each network, indicating whether the subsequent recall effect was greater in the music condition than in the no-music condition. P-values were corrected using false discovery rate to account for multiple comparisons across the 9 networks.

### Supplementary Materials

#### S1: Auditory Cortex RSA

To check that participants were encoding the distinctive features of the six songs that we selected in the scanner, we ran a representational similarity analysis (RSA) on music-condition participants to measure whether neural pattern similarity in auditory cortex was greater for same-song comparisons than different-song comparisons. Auditory cortex is well known to play a critical role in representing the acoustic features of music (Peretz et al., 1994; Zatorre et al., 2002; Norman-Haignere et al., 2022), making it a strong candidate for assessing song-level pattern similarity. For this analysis, we computed representational similarity matrices (RSMs) for each individual music participant by correlating their song-specific event patterns with the group average using a leave-one-out approach; these matrices recorded the average pairwise similarity between all 6 songs, measured using Pearson correlation (for same-song comparisons, we only included comparisons between different instances of the same song). Supplementary Figure 1a shows the average of all of the participant-specific RSMs. A one-tailed (on-> off-diagonal) paired samples t-test was conducted to test whether same-song similarity was greater than between-song similarity. We found that same-song pattern similarity was significantly greater than between-song similarity for the selected songs (*t*(23) = 13.152, p < 0.0001; Supplementary Figure 1b).

**Supplementary Figure 1.**
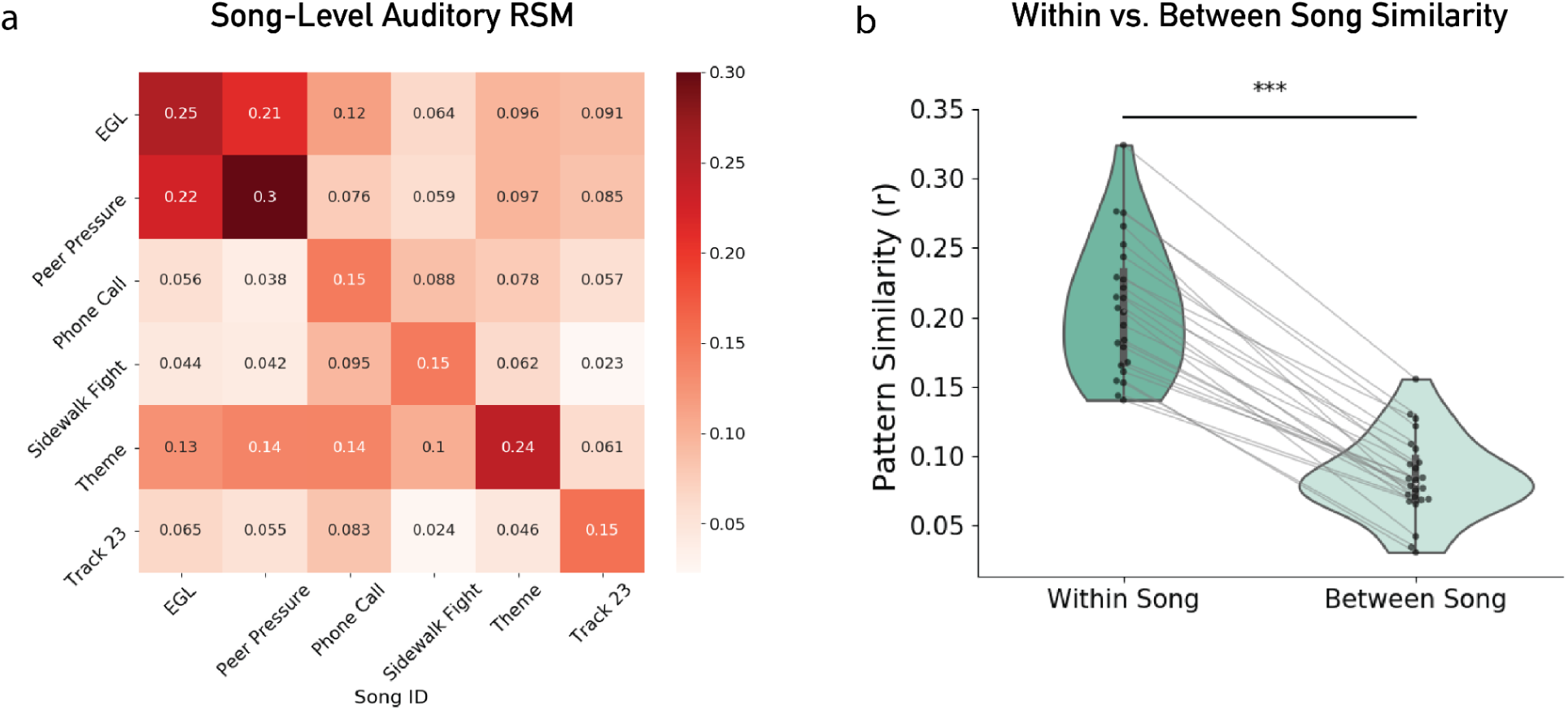
Auditory cortex RSA. Panel a displays the RSM for auditory cortex averaged across all participants in the music condition (n=24). Panel b shows the distributions of *r*-values for same-song comparisons and between-song comparisons (each dot represents a participant). T-test results show that within-song similarity is significantly greater than between-song similarity (*t*(23) = 13.152, p < 0.0001).

#### S2: Remembered vs. Forgotten Reactivation Scores Across All DMN Parcels

We computed reactivation scores for both remembered and forgotten scenes across all DMN parcels (DMNa, DMNb, and DMNc) within the music and no-music conditions. One-tailed paired samples t-tests were performed between remembered and forgotten scores for each ROI (Supplementary Figure 2). When controlling for multiple comparisons within the music condition, ROIs that survive the threshold criterion (q ≤ 0.1) include left IPLa-1 (q = 0.08), left PMC-4 (q = 0.03), and right PMC-2 (q = 0.08). No ROIs survived this criterion in the no-music condition.

**Supplementary Figure 2.**
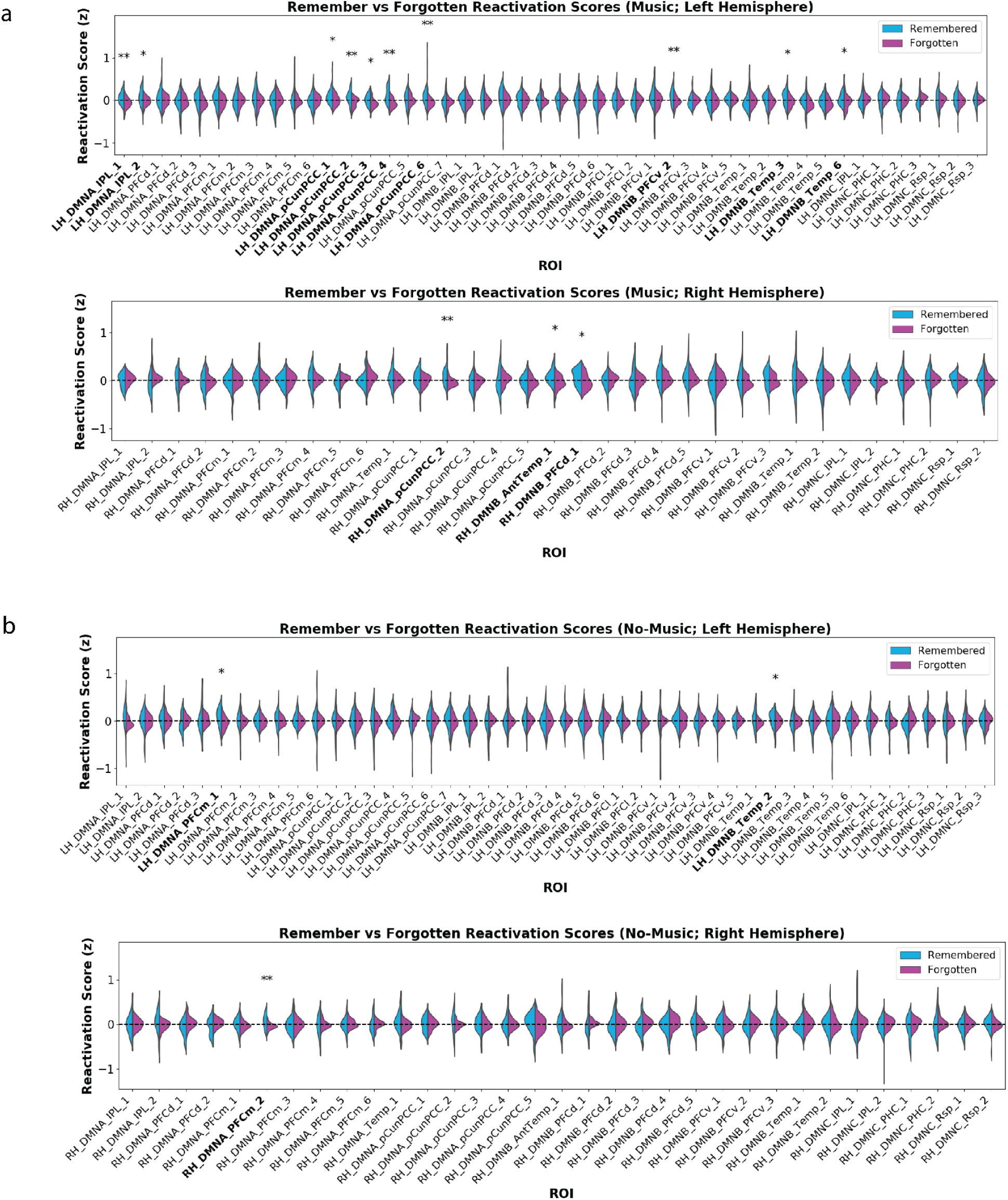
Subsequent recall effect for all DMN ROIs. (a) Shows subsequent recall effects for all DMN ROIs (n=79) within the music condition. The top row shows results for the left hemisphere and the bottom row shows results for the right hemisphere. (b) Shows subsequent recall effects for all DMN ROIs in the no-music condition. Significance levels in this plot were not corrected for multiple comparisons (see S2 text for corrected results). Labels of ROIs with uncorrected significance of p < 0.05 are bolded. Uncorrected significance is also marked with * for p < 0.05 and ** for p < 0.01.

#### S3. Behavioral Recall Performance

##### Participant-level recall performance

We first assessed recall performance at the participant level, computing each participant’s mean recall score and then comparing these scores across the music and no-music conditions. This analysis was conducted for three scene categories: *repeated music scenes* (i.e., the scenes included in the neural reactivation analysis)*, all scenes containing music* (regardless of repetition), and *scenes without music*; comparisons were enacted with independent samples, one-tailed (music > no-music) t-tests. Supplementary Figure 3a visualizes these comparisons, where each black dot represents an individual participant’s mean recall score.

For *repeated music scenes*, the music group exhibited slightly higher mean recall scores than the no-music group, but this difference was not statistically significant (*t*(46) = 0.62, p = 0.27). When considering *all scenes containing music*, we observed a marginally significant trend, with music participants recalling more than no-music participants (*t*(46) = 1.47, p = 0.074). Despite not reaching significance, this trend suggests a potential facilitative effect of music on recall. Moreover, it suggests that this facilitative effect may extend to scenes without repeated music (i.e., the presence of music at encoding may boost recall even if that music is not repeated later).

For *scenes without music,* there was no significant difference between the groups (*t*(46) = 0.3, p = 0.382), though the music group had a numerically higher recall score.

##### Scene-Level Recall Performance

Since the same scenes were viewed in the music and no-music conditions, we conducted a scene-level analysis to assess whether differences in free recall performance were present across the two conditions. For each scene, we calculated the proportions of participants recalling it under the music and no-music conditions, and then we computed the difference in these proportions. Supplementary Figure 3b shows the scene-wise differences in recall proportions (*Music – No-Music*) for all scene types. To assess whether these scene-wise differences were reliably positive across scenes (indicating better recall of scenes in the music condition), we ran a one-sample, one-tailed t-test on the difference scores against zero, with degrees of freedom equal to the number of scenes minus one.

We found a significant recall advantage for music in both *repeated music scenes* (*t*(66) = 2.28, p = 0.013) and *all scenes containing music* (*t*(244) = 6.77, p < 0.0001). However, there was no significant difference for scenes without music (*t*(161) = 1.16, p = 0.123).

We also tested whether scene-level recall differences (music minus no-music) were greater for all music scenes compared to scenes without music, and whether differences for repeated music scenes were greater than those for no-music scenes. We found that scene-level recall differences were significantly greater for all music scenes than no-music scenes (*t*(405) = 3.18, p < 0.001), but this was not the case when we compared repeated music scenes to no-music scenes (*t*(227) = 0.88, p = 0.19).

This scene-level analysis provides a complementary perspective to the participant-level approach. A limitation of this analysis is that it does not permit generalization to new participants. A benefit of this analysis is that each scene is presented in both the music condition and the no-music condition, permitting use of a paired samples t-test that controls for variability in recall performance across scenes (by contrast, we could not use a paired samples t-test in the participant-level analysis shown in Supplementary Figure 3a, since each participant only appears in one condition).

**Supplementary Figure 3:**
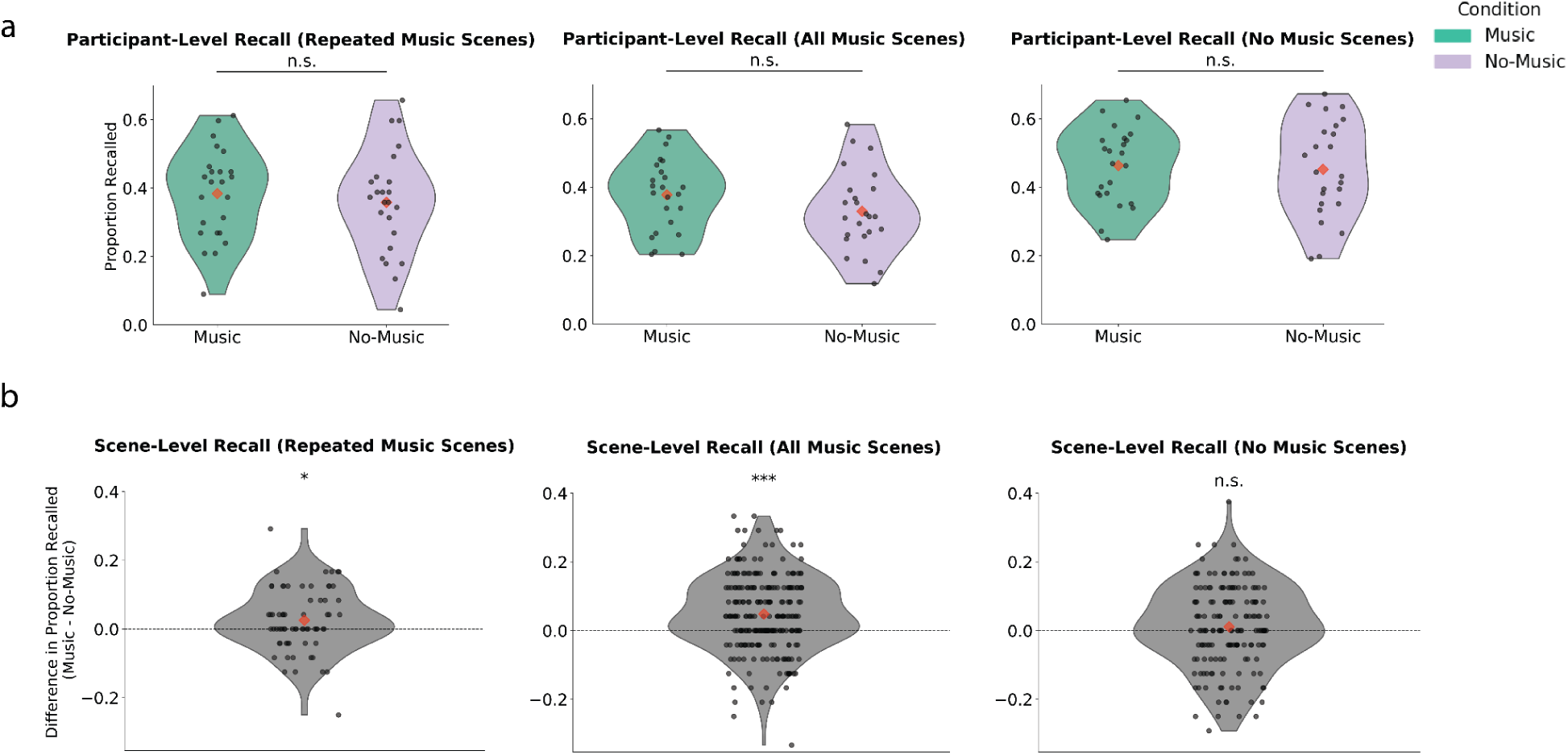
Behavioral recall performance between conditions. (a) *Participant-level recall performance by condition for all scene types*. Violin plots show the distribution of proportion recalled scores across participants for the music and no-music conditions within each scene type. Each dot represents an individual participant and red diamonds indicate the group mean. While the music group showed slightly higher mean recall scores for repeated music scenes, the difference did not reach statistical significance (*t*(46) = 0.62, p = 0.27). The difference was marginally significant for all scenes containing music (*t*(46) = 1.47, p = 0.07); there was no significant difference for scenes that did not contain music (*t*(46) = 0.3, p = 0.38). (b) *Scene-level recall performance for all scene types.* Here, each dot represents a scene and its y-axis coordinate plots the difference in proportion recalled (Music – No Music) for that scene. Results are plotted for all scene types. Proportion recalled was significantly greater in the music condition for repeated music scenes (*t*(66) = 2.28, p = 0.013) and all music scenes (*t*(244) = 6.77, p < 0.0001), but not for scenes that did not contain music (*t*(161) = 1.16, p = 0.123). We also found that scene-level differences in proportion recalled were significantly higher for all music scenes than for scenes without music (*t*(405) = 3.18, p < 0.001); differences in proportion recalled were numerically higher for repeated music scenes than no-music scenes but these differences were not significant (*t*(227) = 0.88, p = 0.19).

#### S4. Reactivation Effects (Not Split by Subsequent Memory)

We tested whether cortical regions exhibited music-evoked reactivation during movie encoding. Analysis steps are the same as those described under *computing reactivation scores* in the *Methods* section, however, reactivation scores were not sorted into bins of remembered or forgotten. Instead significance was tested by evaluating whether subject-level reactivation scores were significantly greater than zero using a one-sample t-test at each ROI. Supplementary Figure 4 shows results when reactivation was tested in a pre-defined set of ROIs (Supplementary Figure 4a) and as a parcel-based searchlight (Supplementary Figure 4b). When performing the analysis using predefined ROIs, no ROIs showed significant evidence of reactivation. However, when performing the analysis as a parcel-based searchlight, significant reactivation was observed in right angular gyrus (*t*(23) = 3.957, q < 0.05) and left parahippocampal cortex (*t*(23) = 2.968, q < 0.1).

**Supplementary Figure 4:**
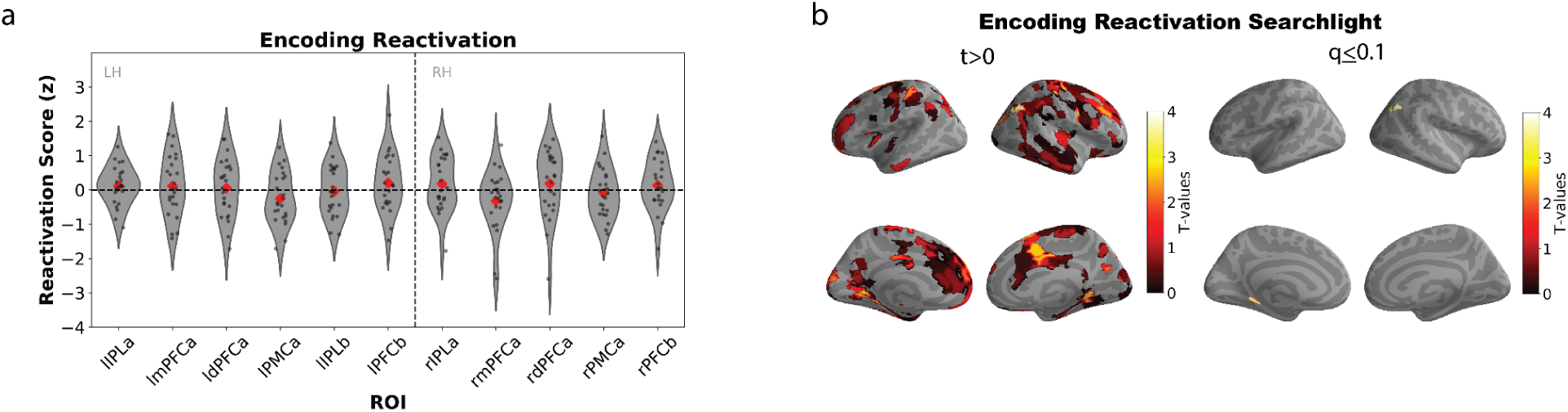
Reactivation effects, not split by subsequent memory. (a) First, we tested whether reactivation occurred in regions of the DMN. When testing for this within pre-defined DMN ROIs, we did not observe significant reactivation in these regions. (b) **Left.** Results of full-brain parcel-based searchlight looking for reactivation (t > 0). **Right.** Significant reactivation was observed in right angular gyrus and left parahippocampal cortex after correcting for multiple comparisons within the DMN (q ≤ 0.1).

#### S5. Reactivation Predicts Subsequent Recall Searchlight: Statistics for Parcels Passing FDR Correction (Visualized in Figure 1D Right Panel)

| Parcel | T-value (df = 23) | Q-value |
| --- | --- | --- |
| Left IPLa-1 | 3.056 | 0.08 |
| Left PMCa-2 | 2.799 | 0.1 |
| Left PMCa-4 | 3.925 | 0.027 |
| Right PMCa-2 | 3.014 | 0.08 |

#### S6. Reactivation Predicts Subsequent Recall (Controlling for ISC) Searchlight: Statistics for Parcels Passing FDR Correction (Visualized in Figure 2F Right Panel)

| Parcel | T-value (df = 23) | Q-value |
| --- | --- | --- |
| Left IPLa-1 | 2.631 | 0.08 |
| Left PMCa-1 | 2.538 | 0.09 |
| Left PMCa-2 | 3.281 | 0.04 |
| Left PMCa-3 | 3.103 | 0.04 |
| Left PMCa-4 | 3.952 | 0.03 |
| Left PMCa-6 | 2.748 | 0.08 |
| Right PMCa-2 | 3.121 | 0.04 |
| Left TEMPb-4 | 3.122 | 0.04 |

## Acknowledgements

Mark Pinsk, Leigh Nystrom, Nicholas DePinto, Kathy Shi, Savannah Born, members of Janice Chen’s lab, James Antony, Sam Zorowitz, Lakshmi Govindarajan, Nina Rouhani, Robert Hawkins, Aaron Bornstein, Rolando Masis, Emily Eyestone, Tankut Can, and members of the Princeton Computational Memory Lab, Hasson Lab, Music Cognition Lab at Princeton, and McDermott labs not mentioned above.

## Funding

Support for this project was provided by F99 NS118740 awarded to JW, and R01 MH112357 awarded to UH and KN.

## Notes

### Competing Interest Statement

The authors have declared no competing interest.

